# A signature of Neanderthal introgression on molecular mechanisms of environmental responses

**DOI:** 10.1101/2021.03.15.435179

**Authors:** Anthony S Findley, Xinjun Zhang, Carly Boye, Yen Lung Lin, Cynthia A Kalita, Luis Barreiro, Kirk E Lohmueller, Roger Pique-Regi, Francesca Luca

## Abstract

Ancient human migrations led to the settlement of population groups in varied environmental contexts worldwide. The extent to which adaptation to local environments has shaped human genetic diversity is a longstanding question in human evolution. Recent studies have suggested that introgression of archaic alleles in the genome of modern humans may have contributed to adaptation to environmental pressures such as pathogen exposure. Functional genomic studies have demonstrated that variation in gene expression across individuals and in response to environmental perturbations is a main mechanism underlying complex trait variation. We considered gene expression response to *in vitro* treatments as a molecular phenotype to identify genes and regulatory variants that may have played an important role in adaptations to local environments. We investigated if Neanderthal introgression in the human genome may contribute to the transcriptional response to environmental perturbations. To this end we used eQTLs for genes differentially expressed in a panel of 52 cellular environments, resulting from 5 cell types and 26 treatments, including hormones, vitamins, drugs, and environmental contaminants. We found that SNPs with introgressed Neanderthal alleles (N-SNPs) disrupt binding of transcription factors important for environmental responses, including ionizing radiation and hypoxia, and for glucose metabolism. We identified an enrichment for N-SNPs among eQTLs for genes differentially expressed in response to 8 treatments, including glucocorticoids, caffeine, and vitamin D. Using Massively Parallel Reporter Assays (MPRA) data, we validated the regulatory function of 21 introgressed Neanderthal variants in the human genome, corresponding to 8 eQTLs regulating 15 genes that respond to environmental perturbations. These findings expand the set of environments where archaic introgression may have contributed to adaptations to local environments in modern humans and provide experimental validation for the regulatory function of introgressed variants.

## Introduction

Studies of ancient DNA samples in recent years have discovered that interbreeding occurred between modern humans and our extinct relatives [1, 2, 3]. Approximately 2% of the genome in modern Europeans and Asians is the result of introgressed Neanderthal sequences [4]. However, Neanderthal ancestry is estimated to be 1.54x the genome average in genomic regions with the lowest density of functional elements. The uneven distribution of these introgressed sequences in the human genome suggests that natural selection may have acted to remove Neanderthal sequences that were deleterious for modern humans [5]. The presence of introgressed archaic sequences in the human genome, instead, raises the question of whether these sequences were maintained in the human genome because they have functional or even beneficial impact in modern humans [6, 7, 8]. Accordingly, Neanderthal introgressed haplotypes have been found to occur at higher frequency than expected by drift and to impact genes with important biological functions for human adaptations in different environments [9, 10]. These include for example BNC2, a gene associated with skin pigmentation [5, 11] and several genes with important immunological functions [12]. However, introgressed non-synonymous variants generally are predicted to have milder functional effects than non-archaic alleles [13].

Gene regulatory variants in the human genome are associated with complex traits and disease risk, supporting the concept that these variants have important phenotypic consequences [14, 15]. Identifying functional non-coding variants that regulate transcriptional processes poses severe challenges because we cannot directly predict function from sequence. Several approaches have been proposed to computationally predict the effect of a nucleotide change on transcription factor binding and gene expression, including methods that are based on sequence content only [16, 17, 18] and methods that integrate sequence content information with functional genomic data, like DNase-seq [19, 20, 21].

Recent studies have focused on understanding the role of Neanderthal introgression on gene regulation in humans using genetic variants which are associated with gene expression (expression quantitative trait loci, or eQTL) from the multi tissue GTEx consortium [22]. Two independent studies used eQTL mapping and allele-specific expression analysis, respectively, to show that a substantial number of Neanderthal introgressed haplotypes harbor regulatory variants in the human genome [13, 23]. Interestingly, brain and testis are the tissues with lower expression of introgressed haplotypes, supporting the hypothesis that Neanderthal introgression may have affected fitness through gene regulation. These observations suggest that it is possible to analyze functional patterns of Neanderthal introgressed sequences to identify potential selection effects on overall pathways or molecular mechanisms.

Studies of genetic regulation of gene expression in cells exposed to pathogens confirmed a role for Neanderthal introgression in immune functions important for the response to bacterial and especially viral infections [24, 25]. Adaptation to pathogen exposure is part of a broader set of signals detected in the human genome and interpreted as adaptations to local environments in human populations. In addition to pathogen exposures, human populations adapted to climate, diet and high altitude [26]. One of the limitations in studies of adaptations to local environments is the difficulty to collect and utilize data on past environmental exposures.

More generally, dissecting the complex environment that we are exposed to today and in the past is a difficult task that requires some necessary simplifications. For example, population genetics studies have summarized past environments with latitude or historical climate and subsistence data [27, 28]; while epidemiological studies generally focus on current measurable lifestyle environments, such as smoking and drinking. An alternative approach to study genetic variants that modulate the response to environmental changes is through *in vitro* systems. Though a simplified version of organismal environments, cultured primary cells exposed to controlled treatments have been pivotal in identifying genetic variants that regulate the response to a variety of different exposures including drugs, hormones, pathogens and other chemical stimuli. We recently demonstrated that this approach can be scaled to investigate hundreds of cellular environments, defined as the combination of specific cell types exposed to different treatments [29]. Approximately 50% of the genes under genetic regulation of transcriptional response are also associated with complex traits variation in humans, thus confirming that the molecular phenotypes measured in this experimental setting (gene expression responses) are important mechanisms in human complex traits variation [29].

Here we used this approach to investigate the role of Neanderthal introgression in modern human response to environmental perturbations. We use a functional annotation of regulatory variants that disrupt transcription factor binding, to predict the regulatory function of Neanderthal introgressed alleles and provide an experimental validation of their regulatory function in human cells.

## Results

### Role of Neanderthal introgressed variants in transcription factor binding and gene regulation

The availability of large amounts of genetic data from human populations and of a variety of different functional annotations makes it possible to explore several unresolved questions on the role of Neanderthal introgression in human history. To analyze the contribution of Neanderthal introgression to modern human response to environmental perturbations, we first refined the definition of introgressed Neanderthal variants. Specifically, we considered variants in previously defined introgressed regions [2] and also present in 2 of 3 high-quality Neanderthal genomes [30, 31, 32], but absent in all African samples from the 1000 Genomes project (Figure S1). We annotated 177,578 SNPs from the 1000 Genomes project [33] as Neanderthal introgressed variants (N-SNPs).

One of the main mechanisms used by human cells to respond to environmental perturbations is through transcriptional changes, which are largely the result of changes in transcription factor binding. To investigate the role of Neanderthal introgression in this cellular response, we focused on N-SNPs that are more likely to have a regulatory function. Specifically, we considered N-SNPs that are in active footprints for transcription factors. To annotate these N-SNPs, we considered the catalog of 1000 Genomes SNPs in active transcription factor footprints generated by [21] using the DNase-seq data from ENCODE and the RoadMap Epigenomics. This catalog annotates 5.8 million SNPs in active transcription factor footprints across 153 tissues and 1,372 motifs. We identified 58,967 N-SNP-motif pairs, corresponding to 27,349 unique N-SNPs in active footprints for a total of 1,255 unique transcription factor motifs. For each transcription factor, we investigated the predicted effect of N-SNPs on binding across all the binding sites in the genome. We found that 19,924 N-SNPs (73%) were computationally predicted to alter transcription factor binding (centiSNPs), which was significantly higher than the 66% of all SNPs which are centiSNPs (*p* < 2.2 × 10^−16^). The Neanderthal allele is predicted to decrease binding in 48% of the cases, increased binding in 44% of cases, and both increased and decreased binding depending on the transcription factor in 8% of cases (Figure 2A). N-SNPs in footprints are more likely to disrupt transcription factor binding (centiSNPs) than non N-SNPs (positive enrichment), with a significant enrichment for 434 transcription factor motifs (BH-adjusted p-value < 0.05; enrichment odds ratios: [1.5, 67.7]). The top 15 enriched motifs (OR>38, Table S1) include factors associated with glucose metabolism and also with response to environmental perturbations. For example, MAFA activates expression of the insulin gene (INS) [34]. Mutations in this gene result in the autosomal dominant disease Insulinomatosis and Diabetes Mellitus [35]. Furthermore, when we considered 37 transcription factors that mediate the response to glucose and/or insulin [36, 37], 16 (43%) were enriched for N-centiSNPs in their footprints genome-wide, suggesting a potential contribution of Neanderthal introgression to modern human metabolism.

**Figure 1:**
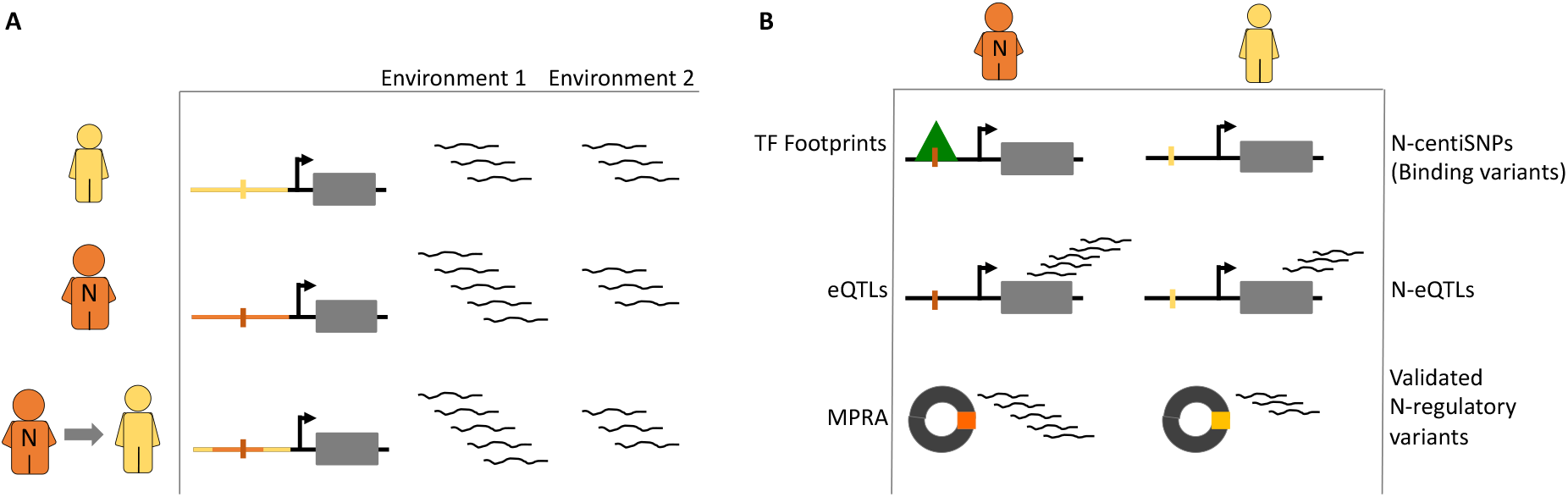
Hypothesis and study design. **(A)** Conceptual framework. We hypothesized that the Neanderthal’s transcriptional response to environmental perturbations was different than modern humans, and introgression of Neanderthal alleles modifies modern human’s response to environmental perturbations. (B) The three approaches used in this study: analysis of Neanderthal introgression in transcription factor footprints, in eQTLs for differentially expressed genes in response to environmental perturbations, and MPRA to experimentally compare the regulatory function of introgressed alleles to modern alleles.

**Figure 2:**
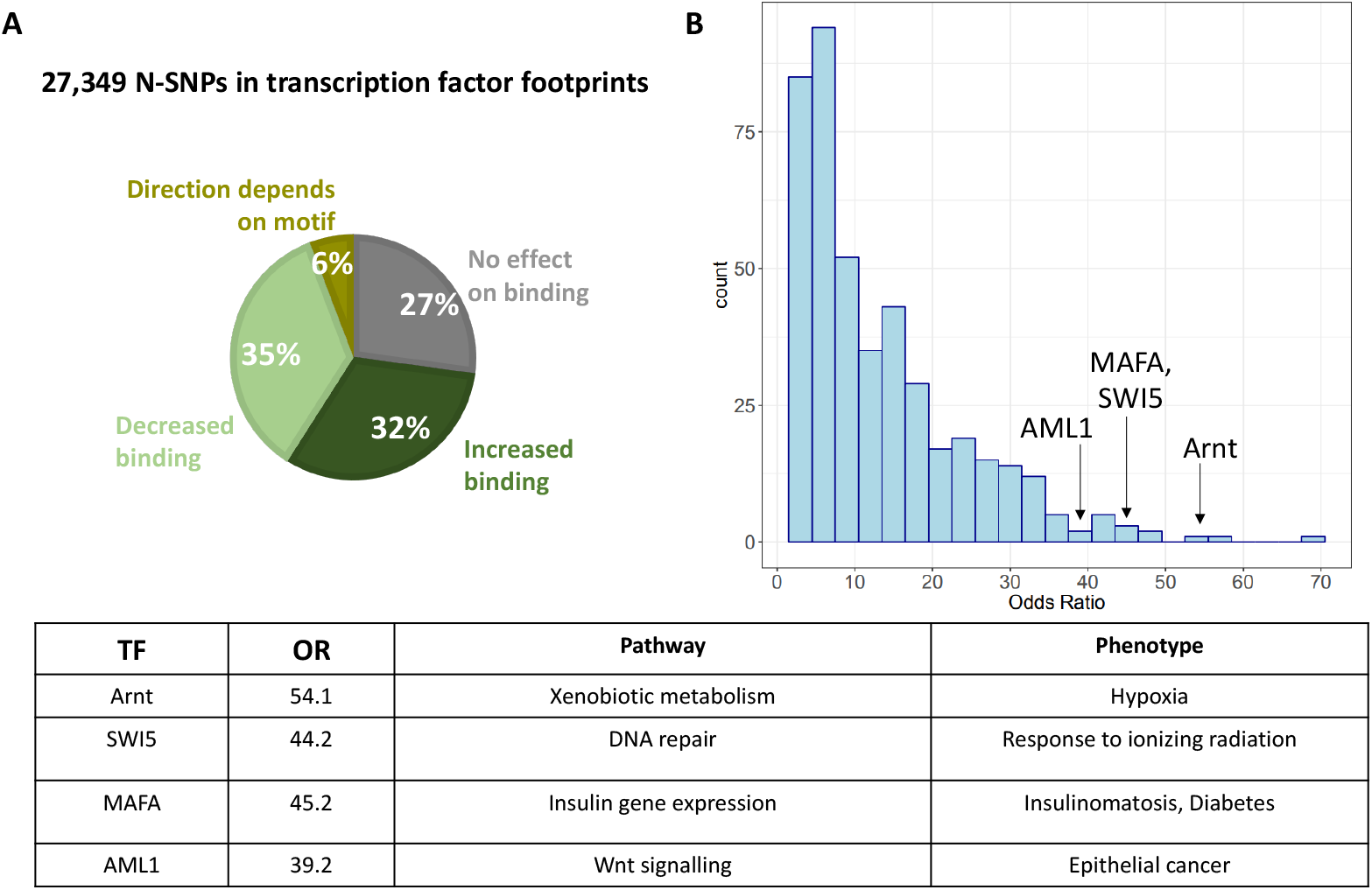
N-SNPs in transcription factor binding sites. **(A)** N-SNPs were annotated in transcription factor binding sites using the centiSNPs annotation, which includes a computational prediction of the SNP’s effect on transcription factor binding. The percentage of N-SNPs in each prediction category is shown in the pie chart. **(B)** Histogram of enrichments of N-SNPs that disrupt transcription factor binding compared to non N-SNPs for 435 transcription factors. **(C)** Phenotypic relevance of the top transcription factor motifs enriched for N-centiSNPs in their binding sites and that are involved in environmental responses.,

Several motifs for transcription factors activated in response to various environmental perturbations were among the most enriched in N-centiSNPs genome-wide. ARNT, which is the most enriched motif, is involved in xenobiotic metabolism and is also a cofactor for hypoxia-inducible factor 1 and NFAT3 (NFATC4) belongs to a family of factors important for T cell activation [38]. SWI5 is a DNA repair protein involved in cellular response to ionizing radiation. Finally, AML1, also known as RUNX1 is involved in hair follicle development and epithelial cancer development. We also found enrichment in footprints for activator protein 1 (AP1), which is known to interact with the glucocorticoid receptor in mediating the response to stress, and for multiple transcription factors (c-Jun, ATF2, and Elk1 [39, 40, 41]) that are downstream of the JNK signaling pathway, which is often induced by environmental stress. These results support previous findings that Neanderthal introgression may play a role in the immune response, and suggest a novel role in regulatory mechanisms for the response to broader environmental stress, and for metabolism.

### Neanderthal introgression regulates the transcriptional response to environmental perturbations

Previous studies have shown that Neanderthal alleles play an important role in human gene regulation [13, 23]. To directly investigate whether Neanderthal alleles regulate the expression of genes that respond to environmental perturbations, we considered expression quantitative trait loci (eQTLs) from the GTEx database and genes differentially expressed in a panel of 52 cellular environments, resulting from 5 cell types and 26 treatments, including hormones, vitamins, drugs, and environmental contaminants [29]. We focused on eQTLs for genes differentially expressed in each cellular environment considered. Overlapping our N-SNPs catalog with the GTEx database, we found that 30,464 GTEx eQTLs (16,065 unique SNPs) in any tissue are N-SNPs. When we considered the set of differential expressed genes in response to treatments, we found that 27,789 N-SNPs are eQTLs for genes differentially expressed in at least one condition, corresponding to 15,026 unique SNPs and 1,629 unique genes (Table S2). For 8 cellular environments we identified a significant enrichment for N-SNP eQTLs in genes differentially expressed, compared to genes that do not respond to the treatment considered (BH-FDR = 0.05). Among these differentially expressed genes regulated by introgressed alleles, we found 239 genes that respond to dexamethasone, a synthetic glucocorticoid that regulates gene expression through activation of the glucocorticoid receptor (GR). The GR is ubiquitously expressed and glucocorticoids regulate different key biological processes depending on the specific body site. For example, in immune cells glucocorticoids act as immuno-suppressors to prevent systemic inflammation and sepsis, while in vascular endothelial cells, glucocorticoids regulate angiogenesis and vascular remodeling. Thus the observed enrichment probably is likely to capture the same signature of Neanderthal introgression on transcriptional regulation of immune response. For example, we found a N-eQTL for the gene RAPGEF3, which encodes for a Ras GTPase. This gene is downregulated in response to dexamethasone in PBMCs and was also among the infection eQTLs identified in monocytes treated with the influenza A virus [25].

We observed a 1.2 fold enrichment for N-SNPs that regulate genes responding to Vitamin D (1,435 genes), which may be linked to adaptive introgression to UV exposure, due to the trade-off between UV exposure and skin pigmentation and its effect on Vitamin D production. For example, BNC2 is downregulated in response to Vitamin D in melanocytes, but upregulated in PBMCs and vascular endothelial cells.

### Adaptive introgression of N-SNPs

Selective pressures on introgressed Neanderthal variation can leave a genomic signature indicative of adaptive introgression. To test for possibility of Neanderthal adaptive introgression at the N-eQTLs in Europeans, we computed summary statistics that have been demonstrated to capture the signature of adaptive introgression, including the RD statistic, the Q95 statistic, and the U20 statistic [8]. The RD statistic is defined as the average ratio of the sequence divergence between an individual from the source population and an individual from the admixed population, and the sequence divergence between an individual from the source population and an individual from the non-admixed population. If a region of the genome is adaptively introgressed into a non-African population, the sequence divergence between Neanderthals and non-African populations should be less than the sequence divergence between Neanderthals and African populations, and, as a result, RD would be small. We identified 6,403 N-SNPs in windows that were significant for the RD statistics (lowest 5% of RD values genome-wide), corresponding to 397 genes (Figure S2A). For each treatment, the number of genes in regions with evidence of adaptive introgression from the RD statistics is proportional to the number of differentially expressed genes regulated by a N-eQTL. The U20 statistic is based on allele frequency in the recipient and donor population and identified adaptive introgression variants based on their elevated allele frequency in the recipient population (> 20%) compared to the donor population. 6,281 SNPs were in regions of adaptive introgression identified via the U20 statistic, corresponding to 296 genes (Figure S2B). On average, 17.9% of differentially expressed genes in response to all treatments are regulated by an N-SNP in a region with a significant U20 statistic. We also computed the Q95 statistics, that considers “high-frequency” archaic alleles, and represents a summary statistic of the site-frequency spectrum in the recipient population. This statistic should be high when a region contains many alleles at especially high frequencies in the recipient population. 6,917 SNPs had a signal of adaptive introgression (highest 5% of Q95 statistic genome-wide) via the Q95 statistic, corresponding to 408 genes (Figure S2C). On average, 25.4% of differentially expressed genes in response to all treatments are regulated by an N-SNP in a region with an adaptive introgression signal via Q95 statistic.

To identify signatures of adaptive introgression, we considered genomic windows that exceed the critical values on all three adaptive introgression summary statistics (the most extreme 5% quantile value from its genome-wide distribution). We therefore identified 1000 N-SNPs in windows that exceed the critical values on all three adaptive introgression summary statistics, hence candidates of adaptive introgression. Of these adaptive introgressed N-SNPs, 384 are N-eQTLs and 363 regulate genes that respond to treatments, for a total of 132 genes(Figure 4A). The proportion of genes regulated by N-eQTLs in regions of adaptive introgression is similar across all treatments (7.5% on average, Figure 4B) and it is not significantly different for genes that do not respond to a treatment.

**Figure 3:**
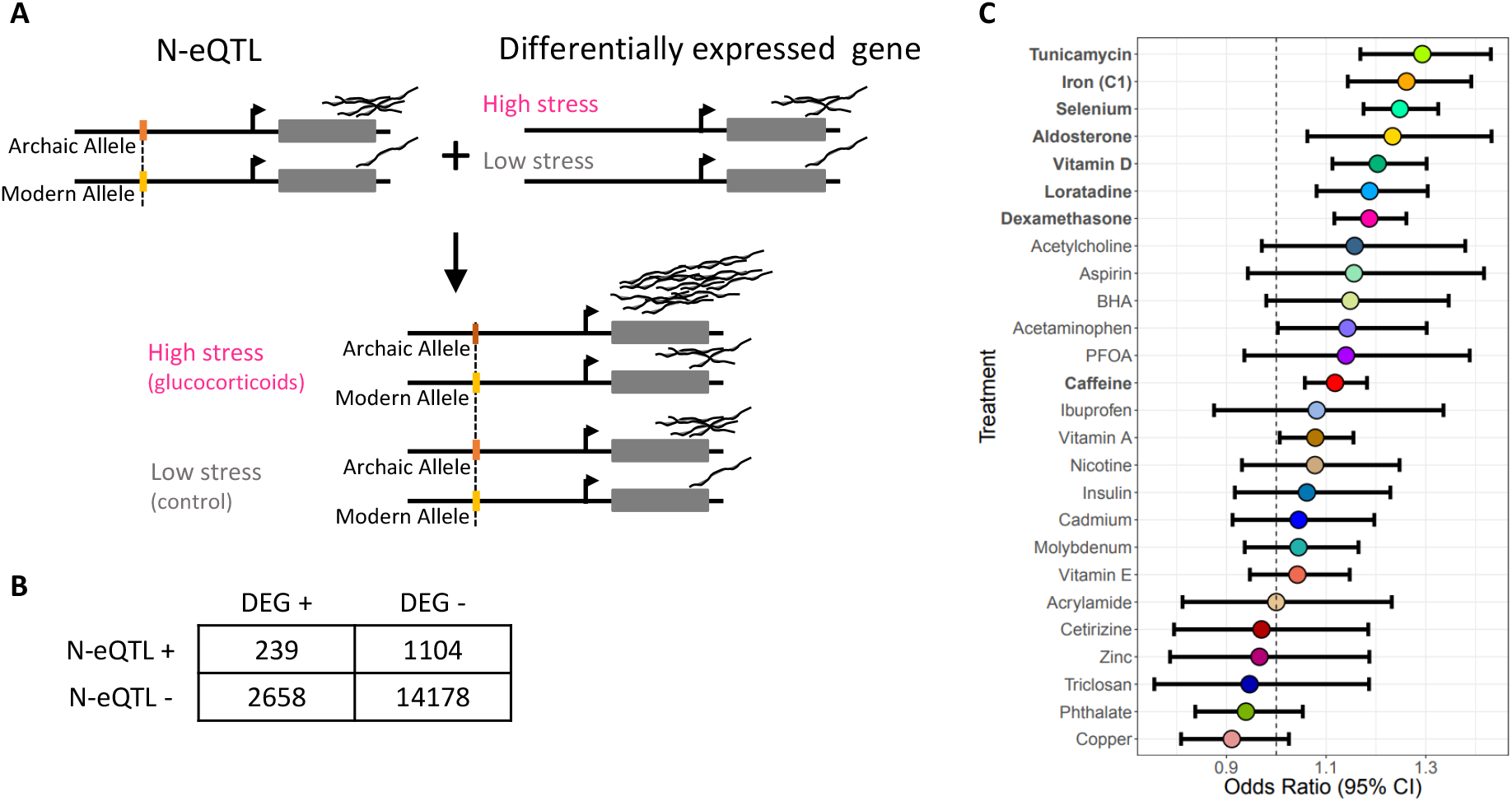
N-SNPs regulate differentially expressed genes. **(A)** Neanderthal introgressed alleles can be linked to the gene they regulate through eQTL mapping signals reported by GTEx. If these genes respond to environmental perturbation, the introgressed variant will contribute to modulating the transcriptional response. **(B)** Example of a contingency table used to test for significant enrichment of N-SNPs for genes that respond to glucocorticoids (dexamethasone treatment). **(C)** Odds ratios and 95% confidence intervals for enrichment of N-SNP eQTLs in genes differentially expressed to each treatment.

**Figure 4:**
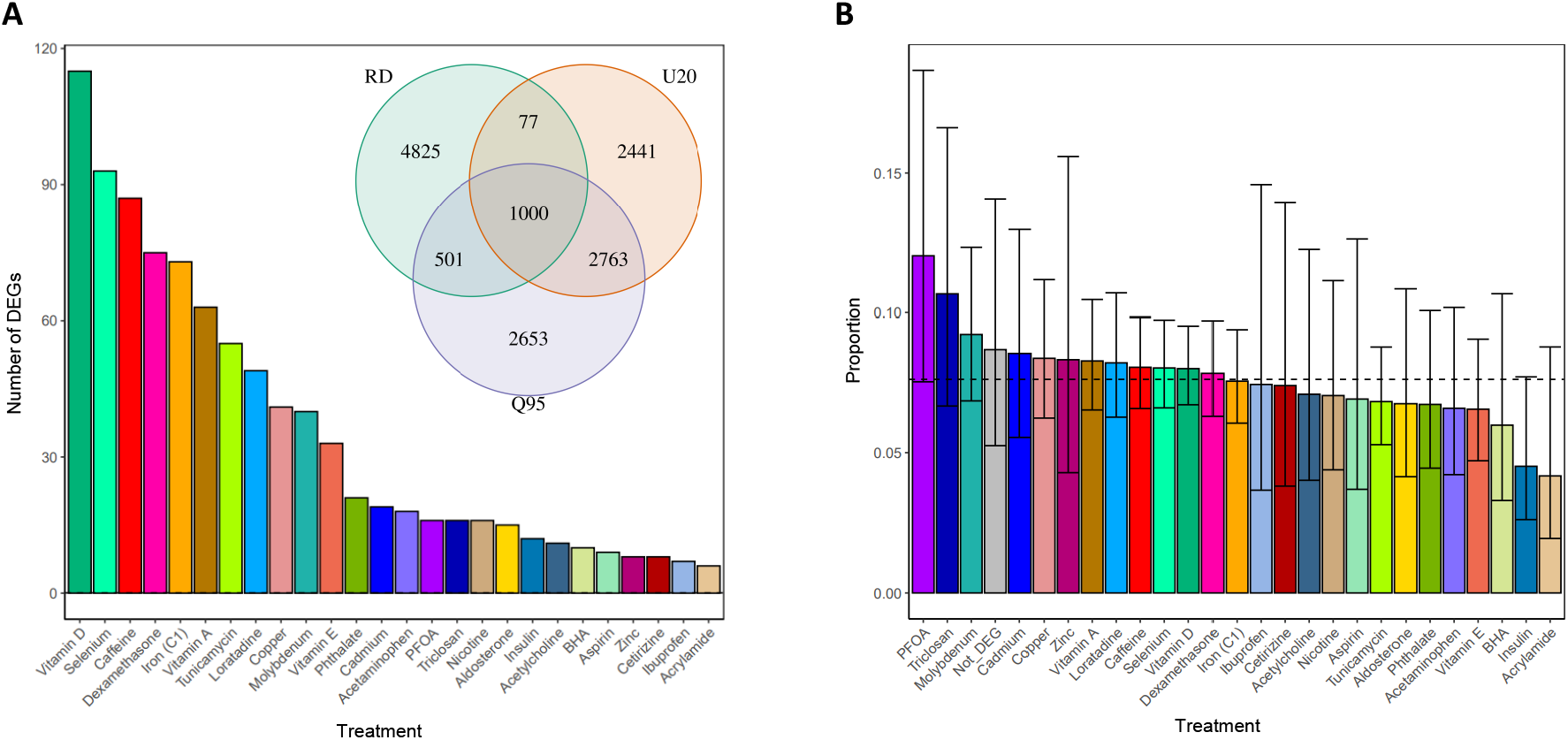
Signals of adaptive introgression in genomic regions containing N-eQTLs. **(A)** Number of genes differentially expressed in response to the treatments and that are regulated by N-eQTLs in regions of adaptive introgression. The inset venn diagram shows the number of N-SNPs which are outliers for each statistic. (**B**) Proportion of genes from A, relative to the number of genes differentially expressed that are regulated by N-eQTLs. The grey bar indicates the proportion of genes regulated by N-eQTLs with adaptive introgression but that are not responding to the treatments.

### Experimental validation of Neanderthal introgressed alleles

Association studies, including eQTLs, are limited in their ability to identify true causal variants in the region of association with the trait of interest. Even considering the lead eQTL for each eGene does not ensure that the true causal variant is being considered. Massively parallel reporter assays (MPRAs) can be used to test the gene regulatory function of DNA sequences carrying individual variants by transfecting them in human cells as part of reporter gene plasmids. To validate N-SNPs that regulate gene expression response, we used BiT-STARR-seq, an MPRA we recently developed to test allele-specific regulatory function for tens of thousands of centiSNPs. We identified 21 N-SNPs (9.3%) with allele-specific regulatory function (BH-FDR<0.1) when considering the 226 N-SNPs included in the library of variants used in [42] (Table S3). This validation rate was expected and was not significantly different than the validation rate in the original study (2,720 out of 43,500, 6.2%). Interestingly, the Neanderthal allele led to increased gene expression for all SNPs. Of these 21 N-SNPs, 13 are centiSNPs, 8 are eQTLs in GTEx and 5 are infection eQTLs in a study of macrophages exposed to listeria and salmonella [24]. The N-centiSNP rs4784812 is in a region of adaptive introgression based on both the RD and the Q95 statistics (RD p-value = 0.046, Q95 p-value = 0.040). The Neanderthal allele at rs4784812 is predicted to alter binding for six transcription factors (NF-Y, CCAAT box, ELK4, or with an ETS DNA binding domain).

We then investigated whether the 8 N-SNPs GTEx eQTLs regulate genes that respond to environmental perturbations. We found that these genes respond to treatments and each of them is differentially expressed in response to several different treatments, as shown in Figure 5B,C. While this set of 8 eQTLs is not a random sample of regulatory N-SNPs, they are likely to regulate genes that are central to the cellular response to environmental perturbations, rather than to specific stimuli.

**Figure 5:**
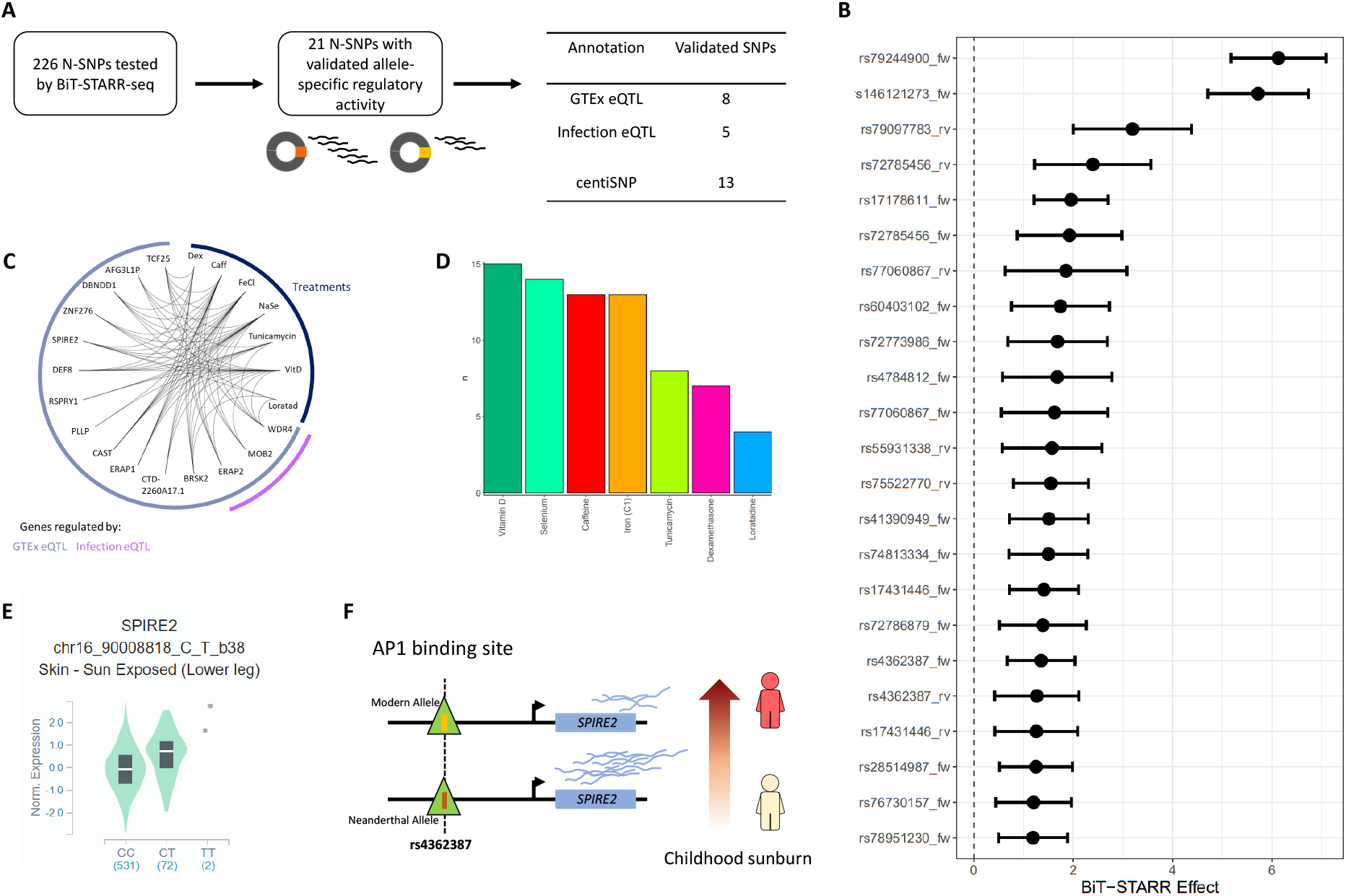
Validation of N-SNP function on gene expression. **(A)** Number of N-SNPs tested and validated by BiT-STARR, with genomic annotations. **(B)** Forest plot showing the effect of the Neanderthal introgressed allele on gene expression in the BiT-STARR-seq experiment. The allelic imbalance is normalized to the allelic ratio in the DNA library. **(C)** Network depicting the genes regulated by N-SNPs validated by BiT-STARR and the treatments in which they are differentially expressed. **(D)** Number of differentially expressed genes per treatment regulated by validated N-SNPs. **(E)** Violin plot of the eQTL signal in the GTEx data for rs4362387 and SPIRE2. **(F)** Graphic representation of a likely mechanism connecting rs4362387 to childhood sunburn based on the molecular signals presented in this study.

Three notable examples are rs72773986, rs28514987 and rs4362387. rs72773986 is an infection eQTL and a GTEx eQTL for ERAP2, and is predicted to alter binding of EPAS1, a key transcription factor for response to hypoxia. ERAP2 encodes a zinc metalloaminopeptidase involved in antigenic response and polymorphisms at this locus are responsible for differential response to bacterial infection and influenza [43]. ERAP2 was also associated with susceptibility to Crohn’s disease [44]. rs28514987 is in a region with significant signals of adaptive introgression by U20 and Q95 statistics (U20 p-value = 0.022, Q95 p-value = 0.031). It is also a GTEx eQTL for BRSK2 and is in a bZIP911 binding site. BRSK2 is expressed in pancreatic islets and negatively regulates insulin secretion [45]. rs4362387 is located within an AP1 binding site and is a GTEx eQTL for SPIRE2 and 6 additional genes in the region. This genomic region also contains the MC1R gene, which is implicated in skin and hair pigmentation and rs4362387 is also an eQTL also for MC1R. The expression of both SPIRE2 and MC1R is highly upregulated by Vitamin D (log_2_ fold change = 2.6 − 2.7, FDR = 0) and likely share regulatory elements. In GTEx, a strong eQTL effect (both in the single tissue and multi-tissue analysis) is observed in the skin tissues, where the T allele (Neanderthal allele) is associated with higher expression of both genes. In the MPRA data, the Neanderthal allele increases expression of the reporter gene, thus showing the same direction of effect as the one observed in the eQTL data, and is also predicted to alter AP1 binding.

**Table 1:**
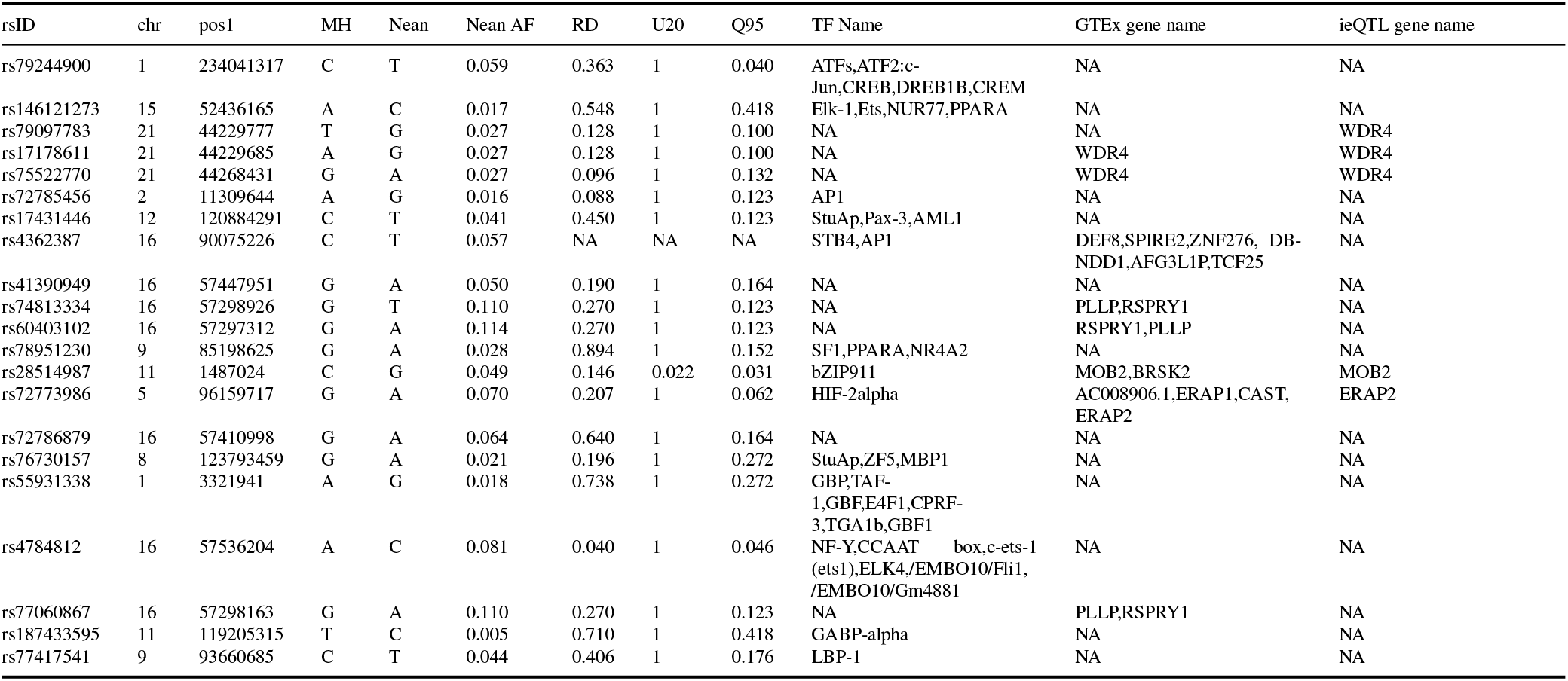
N-SNPs validated by MPRA. Columns 1-12 are: 1) rsID; 2) Chromosome; 3) Position; 4) Modern human allele; 5) Neanderthal allele; 6) Neanderthal allele frequency; 7) RD p-value; 8) U20 p-value; 9) Q95 p-value; 10) transcription factor motif name; 11) GTEx gene name; 12) Infection eQTL gene name

## Discussion

In vitro studies allow researchers to analyze molecular phenotypes in specific environments. While these environments are a simplified version of organismal exposures, they have two key advantages: they are tightly controlled and can simulate environments that are difficult to measure *in vivo*, including ancient environments. To directly investigate whether Neanderthal introgression may contribute to modern humans’ ability to respond to a wider range of stressors, beyond pathogens, we used data on transcriptional response in 35 cellular environments that reflect common individual exposures, including dietary components, environmental contaminants, metal ions, over the counter drugs, and hormonal signaling. While some of these compounds clearly represent very recent exposures in contemporary human history, the underlying hypothesis is that these compounds target existing response mechanisms. For example, endocrine disruptors, such as BPA, which are commonly found in plastics, target the estrogen response pathway [46]. Our approach combines functional genomics and paleogenomics, identifying Neanderthal introgressed variants which may have gene regulatory function. We hypothesized that Neanderthal introgression may have contributed to adaptations to environmental changes by introducing regulatory variants that modify binding of specific transcription factors to a large number of downstream targets. Variation in transcription factor binding sites in humans is more common than non-synonymous variants for specific transcription factors.

This is in line with the idea that changes in the function of master regulators would have large pleiotropic effects[47]. Variation in transcription factor binding sites, instead, introduces specific functional consequences but preserves unaltered the overall function of the transcription factor[48, 49]. Using computational predictions and experimental data on eQTLs and transcriptional response, we assigned a putative function to 33,248 variants which are likely to contribute to modern human response to the environment. For 21 N-eQTLs we validated their ability to modify gene expression in massively parallel reporter gene assays.

While several studies have highlighted a potential contribution of archaic introgression to human phenotypes, the molecular evidence in most cases is suggestive and does not provide experimental data in support of a putative molecular mechanism linking archaic alleles to specific functions. Assigning a function to non-coding variants is a daunting task, which is further complicated by the limited availability of archaic genomes and the inability to directly measure archaic molecular phenotypes [50]. By focusing directly on introgressed alleles and considering fine scale computational predictions, here we have provided an annotation of Neanderthal introgressed alleles at single nucleotide resolution. One advantage of this annotation is that it pinpoints transcription factors whose function is likely to be affected by these archaic variants. For 434 transcription factor motifs Neanderthal introgressed alleles are predicted to modify DNA binding. These putative molecular mechanisms represent ideal candidates for follow-up studies focused on specific transcription factors and/or variants.

Another advantage of focusing on the specific introgressed nucleotides within a full haplotype is the ability to directly test and validate the gene regulatory function of these non-coding variants. Massively parallel reporter assays have been successful in identifying new regulatory variants and validating computational predictions of regulatory elements and variants [51, 52, 53, 54, 55, 56]. One key advantage of this approach is the ability to test tens of thousands of variants in parallel because the assays use a library of plasmid that can be designed to investigate a catalog of variants of interest. At the same time, these assays cannot test genetic variants within their native chromatin context and by design are unable to test long distance interactions. As a consequence, additional introgressed variants that we were not able to validate in this study may indeed regulate gene expression in other environmental contexts. Nevertheless, we provide a proof of concept that applying computational and experimental functional genomics approaches enables researchers to assay the gene regulatory function of archaic alleles in modern human cells. Recently, MPRA data have been used to show that ancestral alleles reintroduced by Neanderthal introgression have activity levels similar to non-introgressed variants, while Neanderthal derived alleles are depleted for regulatory activity compared to reintroduced ancestral alleles [57]. Interestingly, in the dataset we considered, all functionally validated Neanderthal alleles led to increased gene expression. If future studies show this to be the case for other introgressed non-coding variants, it would suggest that variants that disrupt function at the gene regulatory level were not tolerated by modern humans and therefore are less likely to be introgressed in our genomes.

One key result of our study is the finding that Neanderthal introgression may contribute not only to modern human’s ability to respond to pathogens, as previously reported, but likely contributes to our ability to adapt and respond to a broader set of environmental challenges. We discovered that introgressed alleles are enriched in binding sites for transcription factors that regulate response to environmental stressors, including hypoxia and ionizing radiation, and also for transcription factors involved in glucose metabolism. These do not seem to be incidental findings, as they find support in the results from our complementary approaches. For example, we found an enrichment for Neanderthal alleles that disrupt binding for 16 transcription factors involved in the response to glucose and insulin. One of the functionally validated N-eQTLs regulates the expression of BRSK2, a gene that plays a role in the regulation of insulin secretion in response to elevated glucose levels [45]. This N-eQTL is in a region of adaptive introgression, characterized by significant U20 and Q95 statistics.

Several studies in the last decade have identified genetic variants that modify how cells respond to environmental perturbations. These molecular gene-environment interactions have shown that genetic variants have varying effects on gene expression depending on the specific context considered and that these genetic effects are also important for complex trait variation in humans [58, 59, 24, 60, 61, 25, 62, 63, 64, 65, 66, 67, 68, 69]. The majority of these studies have focused on the response to pathogens thus there is limited availability of eQTL datasets to investigate the role of Neanderthal introgression on GxE in modern humans. However, the response to environmental changes depends not only on interaction effects but also on additive effects between environmental variables and genotypes. In other words, an individual carrying a high expression genotype at an eQTL for a gene that responds to a specific environmental stimulus, will have an overall greater response than an individual with a low expression genotype in the same environment. This is because the gene is induced and consequently the genetic effect is amplified [70]. The observed enrichment of N-eQTLs for responding genes suggests a contribution of Neanderthal introgression to modern humans’ transcriptional response to treatments that represent the in vitro counterpart for common exposures, some of which may have represented a selective pressure during human history. Some examples include Vitamin D (a proxy for solar radiation), aldosterone (the main hormone regulating fluid balance), tunicamycin (endoplasmic reticulum stress agent), and glucocorticoids, which exert an anti-inflammatory function in the human body. Glucocorticoids are regulated by the hypothalamus-pituitary adrenal axis in response to stress, thus representing a molecular connection between stress and immune function.

We also found that genes responding to vitamin D, which is produced in response to solar radiation [71], are enriched for Neanderthal eQTLs (OR = 1.2, FDR = 1.9 × 10^−5^). Previous studies suggested that Neanderthal introgressed haplotypes contribute some of the genetic variation underlying the red hair phenotype in modern humans [13, 72]. A non-synonymous variant in the MC1R gene was identified in a Neanderthal genome but wasn’t seen in subsequently sequenced individuals, thus suggesting that it may represent a polymorphic site in Neanderthal populations. The N-eQTL we functionally validated is significantly associated with red hair in the UK Biobank data (*p* = 2.2 × 10^−308^) and is also an eQTL for MC1R and SPIRE, two genes that are differentially expressed in response to Vitamin D. In transcriptome wide association studies, both genes are associated with skin and hair pigmentation phenotypes, including red hair color and childhood sunburn [73] (Figure 5D). Our results raise the possibility that multiple alleles and mechanisms underlie the red hair phenotype not only in humans but also in Neanderthals.

With increasing availability of ancient DNA sequences, combining paleogenomic and functional genomic tools will allow researchers to explore in greatest depth the functional differences and similarities of modern humans from our extinct relatives. Together with other recent studies, we have demonstrated the value of taking these complementary approaches. Our results uncovered a putative role of Neanderthal introgression to modern humans’ ability to adapt and respond to environmental changes, with potential consequences for glucose metabolism and response to solar radiation.

## Methods

### Identification of introgressed Neanderthal variants

We determined a list of putative Neanderthal introgressed SNPs by screening the biallelic SNPs from the 1000 genomes project (1KGP) [33] based on three criteria. The conservative criteria address the notions that Neanderthal introgression occurred outside of Africa and that the Neanderthal-introgressed haplotypes are not observed at certain regions of the human genome. Briefly, the criteria for a Neanderthal-introgressed SNP are: 1) the SNP is within the genomic regions reported to harbor the Neanderthal introgressed haplotypes [2]; 2) the SNP is confidently genotyped in at least two of the three high coverage Neanderthal genomes available [30, 31, 32] and is fixed among the genotyped alleles; 3) the SNP has all of the 504 sub-Saharan Africans (ESN, GWD, LWK, MSL, YRI) in 1KGP fixed for an allele alternative to the Neanderthal one, and is polymorphic among the non-Africans. The rationale of these criteria is to address the notion that much of the Neanderthal introgression occurred outside of Africa. We found a total of 186,871 SNPs that pass these criteria for further analysis. We further removed structural variants and multi-allelic SNPs to end up with 177,578 N-SNPs.

### N-SNPs in transcription factor binding sites

We annotated N-SNPs present in transcription factor binding sites using the centiSNP annotation from Moyerbrailean *et al* [21]. Specifically, we used the centiSNPs Compendium (http://genome.grid.wayne.edu/centisnps/compendium/) which summarizes all 1000 Genomes Project SNPs in 1,372 transcription factor motifs across 153 tissues and estimates the likelihood that a SNP disrupts transcription factor binding. N-SNPs were located within 1,255 transcription factor motifs. For each of the 435 transcription factor motifs with at least 10 N-SNPs within their binding sites, we performed Fisher’s exact test to determine whether N-SNPs in transcription factor binding sites were more likely to disrupt transcription factor binding than non-introgressed SNPs within transcription factor binding sites. Multiple-test correction was performed using the Benjamini-Hochberg procedure [74]. Input data and results from the Fisher’s exact tests can be found in Table S1.

### N-SNP regulation of differentially expressed genes

To link N-SNPs to genes they regulate, we annotated N-SNPs which were significant eQTLs in any of 48 tissues in GTEx v7 (https://storage.googleapis.com/gtex_analysis_v7/single_tissue_eqtl_data/GTEx_Analysis_v7_eQTL.tar.gz). 16,065 N-SNPs were GTEx eQTLs in any tissue, regulating 1,939 genes. We then annotated N-eQTLs which regulated differentially expressed genes in response to environmental perturbation from Moyerbrailean *et al* [29]. We chose all treatment-cell type combinations with at least 1000 differentially expressed genes and downloaded the gene expression data from http://genome.grid.wayne.edu/gxebrowser/Tables/Supplemental_Table_S6.tar.gz. In total, we consider five cell types (melanocytes, peripheral blood mononuclear cells, lymphoblastoid cell lines (LCLs), smooth muscle cells, and human umbilical vein endothelial cells) and the 26 treatments shown in Figure 3B for a total of 52 treatment-cell type combinations.

We identified treatments in which differentially expressed genes were enriched for Neanderthal regulation using Fisher’s exact test. We limited our analysis to genes with an eQTL and tested for an enrichment of N-eQTL genes in genes differentially expressed for each treatment-cell type combination separately. For treatments which were tested in multiple cell types, we meta-analyzed the enrichment odds ratios for each treatment using a fixed effects model with inverse variance weights to obtain a single enrichment per treatment. Multiple-test correction was performed using the Benjamini-Hochberg procedure [74].

### Experimental validation of N-SNPs from BiT-STARR-seq

To validate N-SNPs which cause changes in gene expression, we used data from a massively parallel reporter assay (MPRA) known as BiT-STARR-seq performed in LCLs [42]. Results from BiT-STARR-seq were downloaded from https://genome.cshlp.org/content/suppl/2018/10/17/gr.237354.118.DC1/Supplemental_Table_S1_.txt. We also included an annotation for Neanderthal SNPs which were eQTLs in macrophages infected with *Listeria* or *Salmonella* from Nedelec *et al* [24] (downloaded from https://genome. cshlp.org/content/suppl/2018/10/17/gr.237354.118.DC1/Supplemental_ Table_S4.txt). 226 N-SNPs were tested by BiT-STARR-seq for allele-specific regulatory differences, and 21 were significant after multiple test correction. BiT-STARR-seq results for all 226 N-SNPs can be found in Table S3.

### Adaptive Introgression summary statistics

RD statistic is defined as the average ratio of sequence divergence between an individual from the recipient and an individual from the donor population, and the divergence between an individual from the outgroup and an individual from the donor population. The U20 statistic is defined as the number of uniquely shared alleles between the recipient and donor population that are of frequency <1% in the outgroup, 100% in the donor, and >20% in the recipient population. The Q95 statistic is defined as the 95% quantile of the distribution of derived allele frequencies in the recipient population, that are of frequency <1% in the outgroup and 100% in the donor population [8].

In this analysis, we computed the above statistics in non-overlapping 50-kb windows across the autosomes using modern human genome data from Phase 3 of the 1000 Genomes Project [33]. We used individuals from the CEU population as the recipient population, individuals from YRI population as the non-introgressed outgroup, and the the unphased, high-quality whole genome sequence from the Altai Neanderthal individual [30] as the donor population. We define the statistical values for each eQTL as the AI summary statistic values of each 50kb window that contains the eQTL SNP.

To identify signatures of AI, we adopted an “outlier approach” that we define the critical value for each statistic as the most extreme 5% quantile value from its genome-wide distribution (Table S4). We plotted the genome-wide distribution of the AI statistics as Manhattan plots in Figure S2, with windows containing N-eQTLs highlighted in red. The solid line shows the critical value of each statistic.

## Acknowledgement

Funding to support this research was provided by NIH 1R01GM109215-01 (RPR, FL), NIH F30GM131580 (ASF), and by Wayne State University - Career Development Chair Award (FL). We would like to thank Wayne State University HPC Grid for computational resources and members of the Luca/Pique group for helpful comments and discussions.

## Supplementary Materials

## Supplementary Tables

Table S1: **Enrichment of N-centiSNPs to disrupt transcription factor binding**. Each row represents a transcription factor. Columns 1-12 are: 1) Motif; 2) Odds ratio; 3) Lower confidence interval; 4) Upper confidence interval; 5) Enrichment p-value; 6) Number of N-SNPs which are predicted disrupt transcription factor binding; 7) Number of non-Neanderthal SNPs which are predicted to disrupt transcription factor binding, 8) Number of N-SNPs in transcription factor binding site which are not predicted to disrupt binding, 9) Number of non-Neanderthal SNPs in transcription factor binding site which are not predicted to disrupt binding, 10) FDR of enrichment, 11) transcription factor motif name

http://genome.grid.wayne.edu/Neanderthal/Parse_Fisher.tab

Table S2: **N-eQTLs**. For each N-SNP, we annotated the gene it regulates in GTEx and whether that gene is differentially expressed in Moyerbrailean *et al* [29]. Columns 1-42 are: 1) Chromosome; 2) Position; 3) rsID; 4) GTEx SNP ID. “NA” indicates the SNP is not an eQTL in GTEx; 5) Modern human allele; 6) Neanderthal allele; 7) Neanderthal allele frequency, 8) Ensembl gene id, 9) Gene symbol, 11 - 41) Treatments from Moyerbrailean *et al* [29]. “1” indicates the gene is differentially expressed in that condition in at least one cell type, 42) “1” indicates the gene is not differentially expressed in response to any treatment

http://genome.grid.wayne.edu/Neanderthal/Nean_SNPs.GTEx.DEG.tab

Table S3: **BiT-STARR results for N-SNPs**. Results for the 226 N-SNPs tested by BiT-STARR-seq by Kalita et al [42]. The table was downloaded from Supplemental Table 1 from Kalita et al [42] (https://genome.cshlp.org/content/suppl/2018/10/17/gr.237354.118.DC1/Supplemental_Table_S1_.txt). We added a final indicating the FDR calculated using the 226 N-SNPs only. http://genome.grid.wayne.edu/Neanderthal/Parse_Fisher.tab

Table S4: **Adaptive introgression outliers**.1000 SNPs had outlier values for all three introgression statistics: RD, Q95, and U20. Columns 1-13 are: 1) Chromosome; 2) Position; 3) rsID; 4) GTEx SNP ID. “NA” indicates the SNP is not an eQTL in GTEx; 5) Modern human allele; 6) Neanderthal allele; 7) Neanderthal allele frequency, 8) Ensembl gene id, 9) Gene symbol, 10) 50 kb genomic window where SNP resides, 11) RD statistic, 12) Q95 statistic, 13) U20 statistic http://genome.grid.wayne.edu/Neanderthal/AI_outlier_stats.tab

## Supplementary Figures

**Figure S1:**
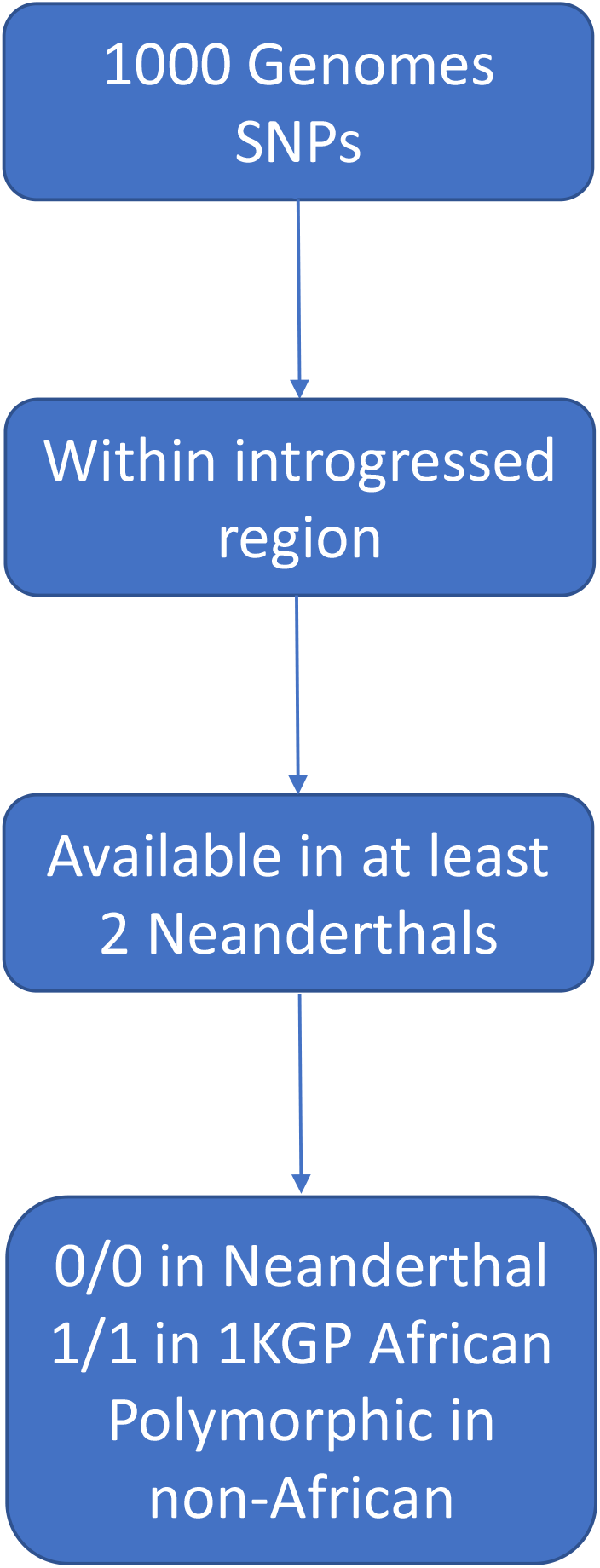
Identification of N-SNPs. Overview of how N-SNPs were selected.

**Figure S2:**
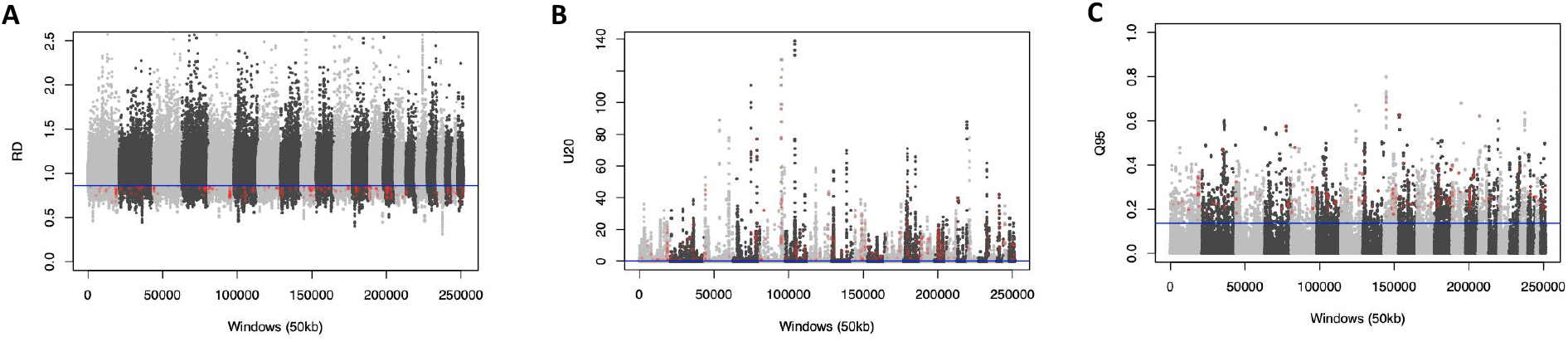
Adaptive introgression statistics. **(A)** RD, **(B)** U20, and **(C)** Q95 statistics for 50kb windows genome-wide. The horizontal line indicates the 5th percentile, and windows with *p* < 0.05 with a N-eQTL are colored red.

